# Panzootic chytrid fungus exploits diverse amphibian host environments through plastic infection strategies

**DOI:** 10.1101/2021.11.29.470466

**Authors:** María Torres-Sánchez, Jennifer Villate, Sarah McGrath-Blaser, Ana V. Longo

**Affiliations:** Department of Biology, University of Florida, Gainesville, FL 32611

## Abstract

While many pathogens are limited to a single host, others can jump from host to host, which likely contributes to the emergence of infectious diseases. Despite this threat to biodiversity, traits associated with overcoming eco-evolutionary barriers to achieve host niche expansions are not well understood. Here, we examined the case of *Batrachochytrium dendrobatidis* (*Bd*), a multi-host pathogen that infects the skin of hundreds of amphibian species worldwide. To uncover functional machinery driving multi-host invasion, we analyzed *Bd* transcriptomic landscapes across 14 amphibian hosts and inferred the origin and evolutionary history of pathogenic genes under a phylogenetic framework comprising 12 other early-divergent zoosporic fungi. Our results not only revealed a conserved basal genetic machinery, but also highlighted the ability of *Bd* to display plastic infection strategies when challenged under suboptimal host environments. We found that genes related to amphibian skin exploitation have arisen mainly via gene duplications. We argue that plastic gene expression can drive variation in *Bd* lifecycles with different mode and tempo of development. Our findings support the idea that host skin environments exert contrasting selective pressures, such that gene expression plasticity constitutes one of the evolutionary keys leading to the success of this panzootic multi-host pathogen.

## Introduction

Multi-host pathogens represent an exceptional case of species evolution in changing environments (1). These host-shifting pathogens can colonize and proliferate in different hosts while evolving adaptations, plausibly, through the interplay between virulence and transmission (2,3). The process of host colonization can drive pathogen diversification if host alternation does not constrain pathogen adaptation to new host environments. When host alternation is prevalent, pathogens might display the same infection strategy across hosts, becoming what has been called ‘jack of all trades and masters of none’. At the same time, multi-host pathogens are resource specialists and can be masters of some or, even, all their hosts (4,5). Accordingly, to exploit their specific resources in both optimal and suboptimal host environments, pathogens can develop either a compromised or a plastic infection strategy to optimize their fitness (6). Characterizing pathogen evolutionary dynamics inducing different infection strategies is essential to understand host shifting success and to predict the pathogen’s ability to colonize new host environments (7). This ability can ultimately contribute to curbing emerging infectious diseases in wild populations (8).

Outbreaks of multi-host pathogens in amphibians, snakes, and bats demonstrate that the ability to exploit novel host environments can occur rapidly with dramatic consequences to naïve hosts (9). One of the best examples is amphibian chytridiomycosis, an emerging infectious disease caused by the batrachochytrid fungi *Batrachochytrium dendrobatidis* (hereafter *Bd*) (10) and *B. salamandrivorans* (*Bsal*) (11). Of particular importance is the hypervirulent *Bd* Global Panzootic Lineage (*Bd*GPL) that emerged at the beginning of the 20^th^ century causing extinctions and severe declines in hundreds of amphibian species worldwide (12–15). In contrast to the non-pathogenic species of the early-diverging phylum Chytridiomycota, the batrachochytrids are waterborne zoosporic pathogens that exploit vertebrate substrates targeting keratinized areas in the amphibian skin (16). Batrachochytrid infection in amphibians arose as a novel trait with the evolution of particular genetic features, including several types of proteases, chitin-binding proteins, and Crinkler effectors (17–21), which are expressed during host exploitation (22). While developing in amphibian skin, *Bd* exhibits an archetypal chytrid lifecycle: flagellated spores encyst, penetrate the substrate through rhizoids, maturate to thalli, and asexually reproduce into zoosporangium (16). This process causes systemic effects that can alter amphibian skin function and structure with lethal outcomes for susceptible hosts (16). In less susceptible species, amphibian hosts rely on multiple strategies to avoid or control disease progression, including skin microbiome interactions, immune system activation, and behavioral traits to reduce exposure (see review (23)). Not surprisingly, *Bd* infection outcome is highly variable across amphibian species and both biotic and abiotic context-dependent (24).

To understand *Bd* pathogenicity and evolution, several studies have explored its population dynamics (25,26), genetic diversity and innovations (12,17–21,27), and gene expression of both *Bd* and amphibian hosts (21,22,28–35). The scope of these studies has been limited to genomic comparisons with few species of the phylum Chytridiomycota (17,21), whereas *Bd* functional machinery has been examined to understand host exploitation only in two species with similar infection outcomes (22). Therefore, it remains unclear whether Bd can exhibit functional plasticity to overcome different host environments. We hypothesized that host species represent different environments, which carry a specific skin microbiome and defense strategies (23), exerting contrasting selective pressures to *Bd*. Hence, each amphibian host is a unique ecosystem with its own environmental conditions due to its specific life history. Hence, we could expect that *Bd* has encountered new ecological opportunities through the exploration and colonization of different amphibian skins. Under these ecological and evolutionary scenarios (i.e., each amphibian species), *Bd* could display a conserved infection strategy showing resilience during host alternation in inadequate/suboptimal host environments, or exhibit trait variation to potentially enhance its fitness across different hosts (i.e., phenotypic plasticity). Due to the extreme success of *Bd* able to infect hundreds of amphibian species, we predicted that *Bd* can mount different/plastic infection strategies depending on the host environment through variation of its functional machinery (i.e., gene expression; Fig. 1A). This predicted plastic infection strategy would precede adaptation and might have been one of the evolutionary keys to the emergence of this panzootic multi-host pathogen.

**Figure 1.**
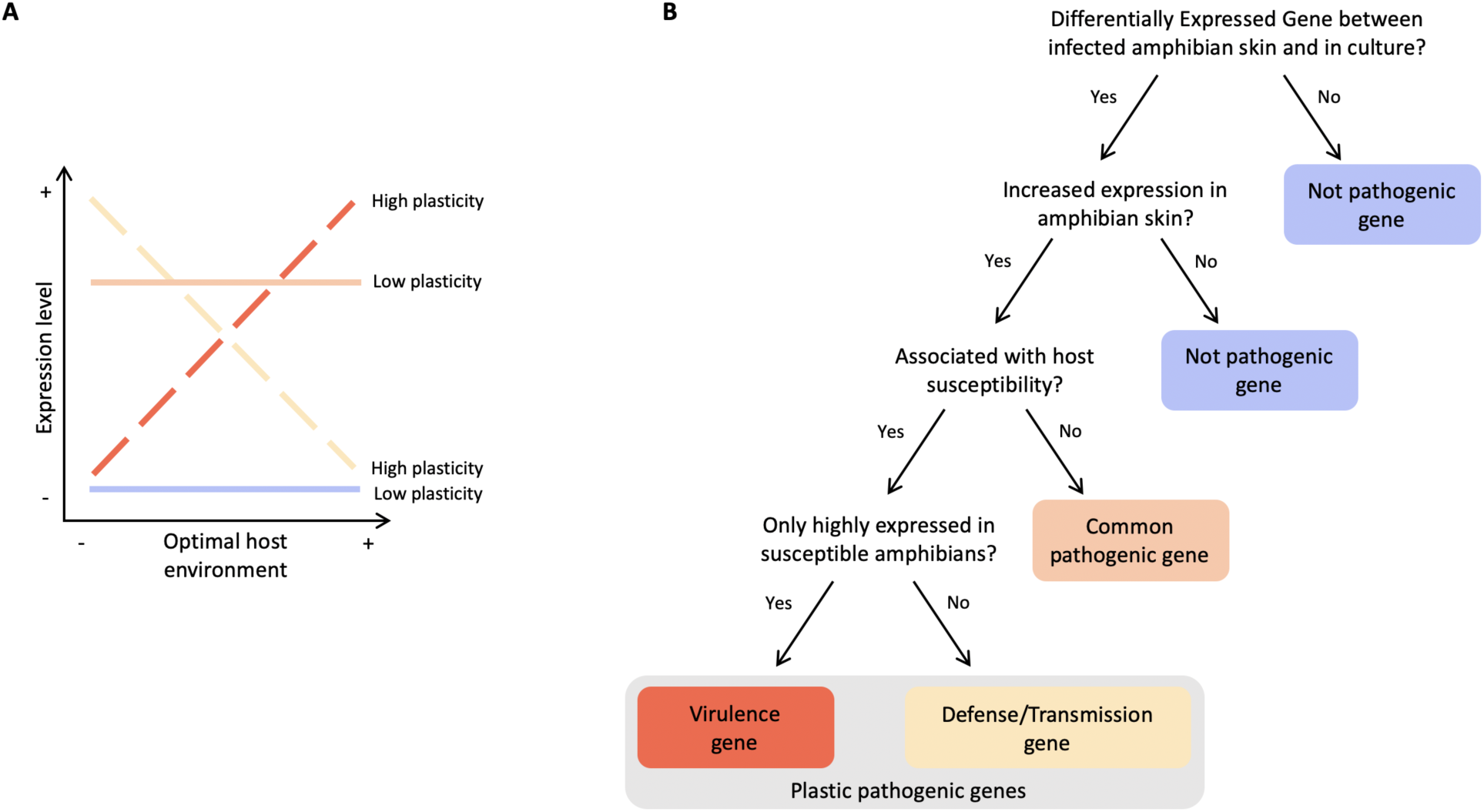
Theoretical gene expression changes driven by host environments. **A**. Schematic illustration of hypothetical gene groups and their expression patterns across different hosts species (form optimal to suboptimal hosts). **B**. Decision tree used to identify candidate pathogenic genes expressed during amphibian host exploitation.

Here, we leverage publicly available transcriptomic datasets (21,22,28–35) and new data from experimental infection challenges (36,37) to investigate *Bd*’s functional machinery across 14 different hosts species in comparison to its expression in culture samples (21,22,31,33). All the animals used in the different studies were experimentally infected with a *BdGPL* isolate, and most individuals were captive-bred or *Bd*-naïve to the strain used in the experimental trials (information about Bd isolates available in Table S1). Across all the studies, infected skin tissues were harvested after necessary time to complete *Bd* life cycle. Hence, these skin expression profiles allowed us to assess the genetic machinery that *Bd* uses to infect a wide range of different host species before adaptive or co-evolutionary processes. Our analyses identified sets of pathogenic genes (i.e. differentially expressed *Bd* genes upregulated in the amphibian skin; see Fig. 1B for tree decision behind our classification) used to successfully proliferate under different host environments. To trace if these genes represent evolutionary novelties, we compared the genes in a phylogenetic framework comprising 12 other early-divergent zoosporic fungi and inferred their origin and evolutionary history. We predicted that *Bd* will express a different genetic machinery in relation to host infection outcomes, in which susceptible hosts provide optimal environments whereas less susceptible hosts (i.e., partially-susceptible and tolerant hosts) lend suboptimal environments for *Bd* proliferation.

## Results

### Bd expressed distinctive genetic machinery across host environments

We recovered 145 *Bd* gene expression profiles from transcriptome samples of the skin of experimentally infected amphibians (112 *in vivo* samples) and from in culture studies (33 *in vitro* samples). Across samples, the maximum number of expressed genes was 8,236 (94.67% of the total genes reported in the *Bd* JAM81 representative genome). The mean number of expressed genes was 7,866 in culture samples, 6,241 in susceptible host samples, 1,060 in partially susceptible host samples, and 858 in tolerant host samples (see Table S1 for expressed gene number per sample). After classifying samples based on the similarity of their gene expression profiles, we found that the first principal component (PC1) explained a high proportion of the total gene expression variance (70.87%, see Table S2 for variance proportion information of the subsequent PCs) revealing several biological patterns (Fig. 2) and no grouping related to technical characteristics, such as sequencing coverage (see Fig. S1) or *Bd* strain (see Fig. S2). We uncovered a similar general expression pattern among *Bd* samples grown *in vitro*, which included two different *Bd* lineages (*Bd*GPL and *Bd*Brazil) (33), as these points clustered at one end of the PC1 (Fig. 2). In contrast, *Bd* gene expression *in vivo* showed greater variability, which was mainly explained by a quantitative gradient related to host susceptibility: expression profiles from susceptible host samples were closer to the profiles from *Bd* isolates, whereas profiles from less susceptible hosts were more distant to these *in vitro* samples (Fig. 2). *Bd* showed greater gene expression variation in the skin of two species (*Plethodon cinereus* and *Litoria verreauxii alpine*), reflecting the experimental design of those studies (29,32). Despite this, we found that PC1 loadings were repeatable, demonstrating that *Bd* expression profiles were conserved across species (*R* = 0.808, CI = [0.608, 0.893], likelihood-ratio-test P = 1.27e-48). We also identified a significant association between species mean PC1 loadings and host susceptibility category (F value = 17.59, df_effect_ = 2 and df_error_ = 11, p-value = 0.000374). Pairwise comparisons indicated that in susceptible hosts *Bd* displayed a different gene expression pattern from both partially-susceptible (Fig. S3, diff = 0.0728, CI = [0.0328, 0.1128], p-value = 0.0012) and tolerant hosts (diff = 0.0731, CI = [0.0354, 0.11009], p-value = 0.0007) while in tolerant and partially-susceptible hosts, *Bd* expressed similar general patterns (diff = 0.0003, CI = [-0.0403, 0.0396], p-value = 0.9997). These differences were decoupled from the evolutionary history of the hosts, as we did not find a phylogenetic signal using the mean PC1 loading across amphibian hosts (Blomberg’s κ = 0.383, p-value = 0.284; Pagel’s λ = 6.6107e-05, p-value = 1).

**Figure 2.**
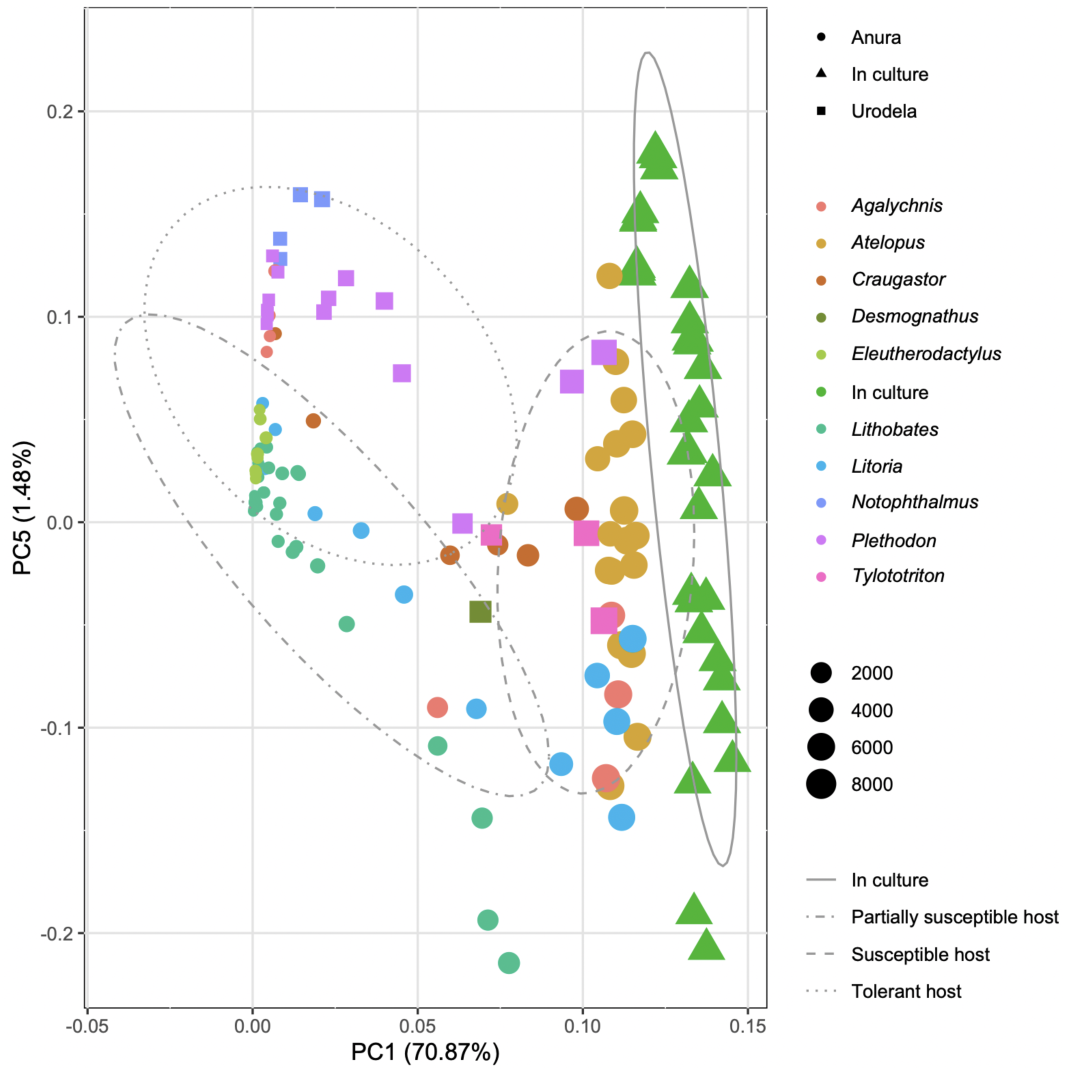
*Bd* gene expression profiles across 14 amphibian hosts. Principal Component Analysis of *Bd* gene expression profiles for 145 samples of skin and cultures. Object shapes and colors differentiate the origin of the samples by amphibian order (Anura (•) and Urodela (◼)) and in culture (▲) while object size illustrates number of expressed genes. Samples are also grouped by ellipses related to host susceptibility.

### Characterization of the common pathogenic Bd machinery

The colonization of amphibian skin, as opposed to the growth in culture, involved a reduction in expression for a large number of *Bd* genes (downregulated genes; Fig. 3A). In tolerant hosts, *Bd* expression profiles showed the greatest percent of downregulation (66.86%), followed by partially-susceptible hosts (60.06%). However, only 17.23% of *Bd* genes were downregulated in susceptible hosts. We identified sets of genes with increased expression (upregulated) in the infected amphibian skin compared to the profiles obtained from cultures (1,033 genes with one or more than one positive unit of logarithmic fold expression change: 577 in susceptible hosts, 819 in partially susceptible hosts, and 564 in tolerant hosts; Fig. 3B and File S1 for detailed information about upregulated genes, including NCBI accession numbers). We defined these genes with increased expression in the amphibian skin as pathogenic genes (Fig. 1B). We detected a core of 300 *Bd* genes that were highly expressed in all the amphibian hosts regardless of their susceptibility category, plausibly involved in host exploitation (Fig. 3B). In this common pathogenic genetic core, we found a battery of genes encoding endopeptidases and carboxypeptidases with extracellular locations or destinations (e.g., aspartic-type endopeptidases, mainly beta-secretases, serine-type endopeptidases, metalloendopeptidases, cysteine-type endopeptidases, metallocarboxypeptidases, and serine-type carboxypeptidases; see File S1). Along with these peptidases, other essential members of *Bd*’s secretome included lipases, a circumsporozoite protein, a beta-lactamase-like protein, Crinkler effector proteins, and regulatory or interference molecules (small delta antigen and replication protein 1a). We also identified genes encoding structural constituents involved in the integrity of the cell wall (such as the pathogen-related yeast protein) and interactive molecules of the cellular membrane including adhesion molecules (halomucin, galectin-3, neurofascin, protocadherin, envelope glycoprotein), voltage-gated channels (sodium, calcium, hydrogen channel proteins) and transmembrane transporters (arginine exporter proteins, methylthioribose transporters, proton pumps, ATP-binding cassette transporters, agnestins efflux protein, high affinity iron permeases). We detected numerous genes encoding chitin deacetylases, several nucleases, and deubiquitinating proteins contributing to the pathogenic genetic machinery of *Bd* with a common increased expression across amphibian hosts.

**Figure 3.**
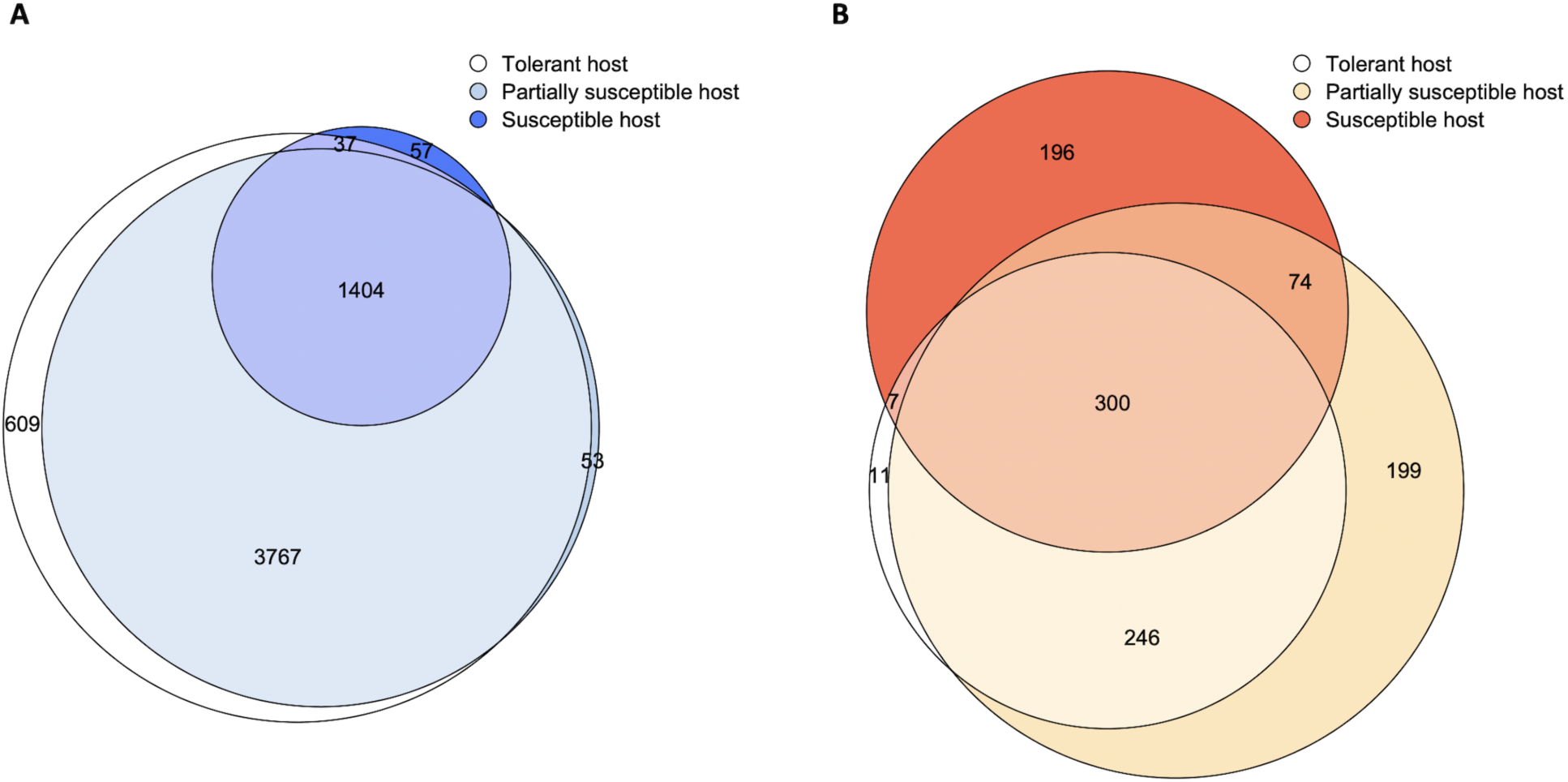
Differential expressed *Bd* genes. Euler diagram showing the numbers of upregulated (**A**) and downregulated (**B**) genes in amphibian skin across host categories.

**Figure 4.**
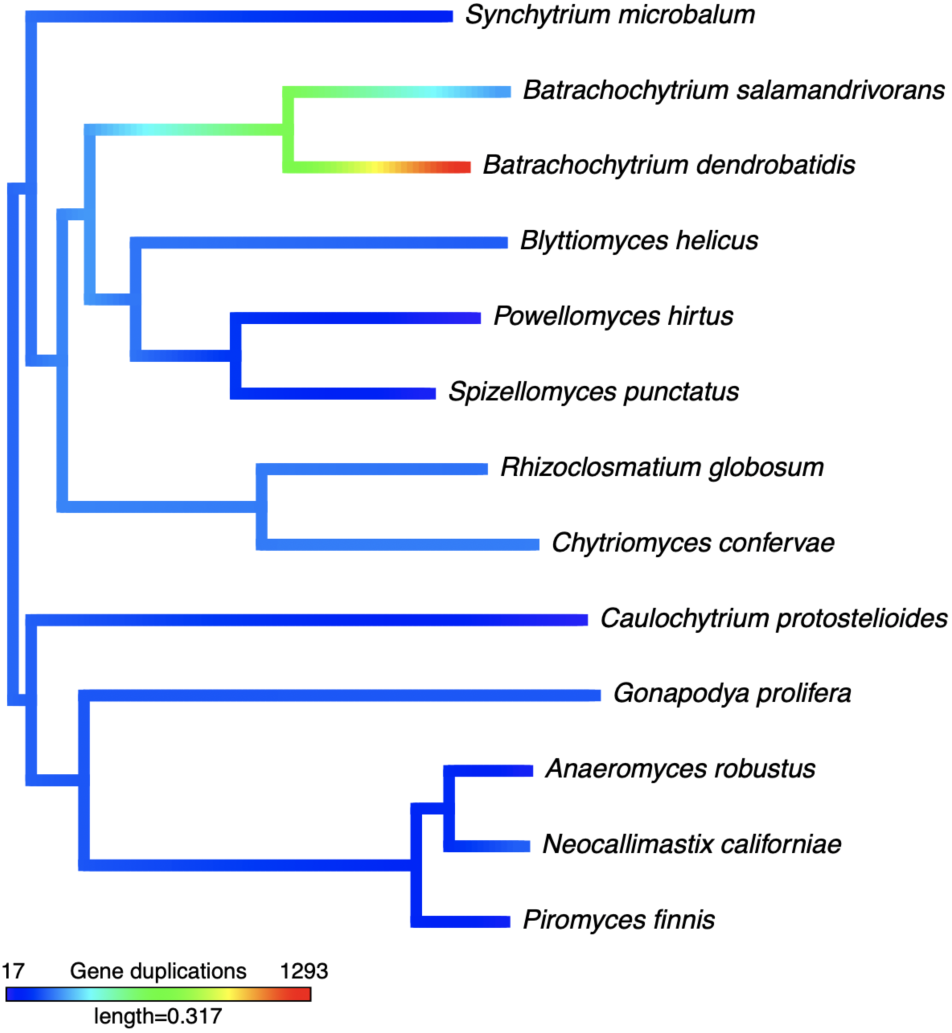
Molecular evolution of *Bd* pathogenic genes. Phylogenetic tree of 13 early-divergent zoosporic fungi showing duplications for the gene families containing the homologs of the *Bd* upregulated genes in amphibian skin (pathogenic genes).

### Sorting Bd pathogenic genes by host environment

From the upregulated *Bd* genes in amphibian hosts, we identified plastic genes that modulated their expression in association with particular hosts categories (see Fig. 1B, 3B, and File S1). We found 196 candidate virulence genes that were highly expressed solely in the susceptible hosts (Fig. 3B). In contrast, we identified 456 putative defense/transmission genes upregulated exclusively in partially-susceptible and/or tolerant hosts. Upregulated genes in susceptible hosts (compared to both partially susceptible and tolerant hosts) were more frequently associated to genes found in sporangia specific samples (χ^2^ = 52.402, p-value = 4.522e-13, Cramér’s V Phi = 0.2892). In addition to the enhanced expression of extra peptidases and transmembrane transporters, *Bd*’s ability to exploit susceptible hosts was characterized by genes related to cell adhesion, predominantly through actin and myosin filaments including anchored proteins to leukocyte cells. Likewise, we detected three chitin-binding genes and a chitinase among the *Bd* genes with adhesion function and high expression in susceptible amphibians. Among these candidate virulence genes, we also uncovered proteins of the Velvet complex, which are conserved regulatory proteins in the kingdom fungi. In contrast, in less susceptible hosts, *Bd* highly expressed additional genes involved in the ubiquitination system, including genes related to the negative regulation of seed maturation and germination with putative defense/transmission functions. Other *Bd* protective mechanisms to fight host defenses could involve toxin production (e.g., gene annotated as pelamitoxin with increased expression in partially susceptible and tolerant hosts). Our results also highlight a large number of genes expressed in less susceptible hosts related to gene silencing, mainly endoribonucleases and interference RNAs or RNAi (six genes Dicer-like protein, two cyclin kinases, an RNA-dependent RNA polymerase, a C6 finger domain transcription factor nscR, and a cross-pathway control protein 1), which could be involved in a bidirectional trans-kingdom communication system.

### Origin and evolution of Bd pathogenic genes

We identified a total of 5,807 candidate gene families (orthogroups) with one or more *Bd* genes across our early-diverging fungi phylogeny that included the closest amphibian pathogenic chytrid fungus (*Bsal*), being 136 of those classified as *Bd* species-specific families. This number represents 36.1% of the total identified gene families across all the studied fungi (Fig. S4) and included 8,096 of 8,700 *Bd* genes. Our results showed 1,602 terminal *Bd* gene duplications (80.71% of duplications in gene families with upregulated genes, see Fig. 3A) and 604 unassigned or orphan genes acquired during the *Bd* evolutionary history. *Bd* pathogenic genes (genes with increased expression in amphibian skin, Fig. 1B) were distributed between orphan genes and gene families (104 unassigned genes and 929 genes in 390 gene families, being 77 of them *Bd* species-specific, File S1). We identified different mechanisms of genetic innovations that might have given rise to novel *Bd* pathogenic genes including the onset of *de novo* genes, horizontal gene transfers (HGT), and duplications of existing genes followed by neofunctionalization. Alternatively, these gene families and/or orphan genes might have arisen from gene losses across this phylogeny and homologs might exist in unsampled species. Among these *Bd* specific gene families, we found several families encoding capsid proteins and envelope glycoproteins that could facilitate cell host invasion as well as *Bd* protection inside the hosts. Specific Crinkler effector proteins related to host interference were also grouped in a *Bd* specific gene family. To respond to interactions with the host environments more efficiently, *Bd* acquired species-specific families that encode serine/threonine-protein kinases, beta-secretases and arginine exporters during its evolution. We found that these genes have highest homology with the arginine transmembrane transporter in the *Serratia* bacteria, which could be the potential donor of a single HGT event followed by several gene duplications. Other candidate HGT events with potential important defense functions were two unassigned *Bd* genes encoding a beta-lactamase and a costunolide synthase. We uncovered other genetic novelties through the expansion of 79 gene families with homolog sequences in other early-diverging zoosporic fungi (14 families with gene members exclusively found on *Bd* and *Bsal*). Plastic pathogenic genes with increased expression in less susceptible hosts were more often classified under *de novo* genes and HGT. Finally, while some gene families had members with high sequence variability, most of the gene families including *Bd* pathogenic genes had sequences under the same evolutionary pressures with a common inferred dN/dS ratio (non-synonymous/synonymous substitutions ratio) among all fungi species (File S1 and Fig. S5). We detected sites with signatures of positive selection in the *Bd* branch (foreground branch) in only five gene families: a gene family annotated as methylthioribose transporters under expansion in *Bd* (positive selection signature in three independent branches leading to *Bd* gene duplications, Fig. 3C) and increased expression in all amphibian species; two families which *Bd* genes had an increased expression in susceptible amphibians (ATP-dependent helicase and multidrug resistance-associated protein); and another two families which *Bd* genes were highly expressed in less susceptible hosts (ATP-synthase subunit beta and elongation factor).

## Discussion

Chytridiomycosis threatens amphibian populations worldwide, however, disease risk varies greatly depending on a myriad of factors, including *Bd* lineage (12,13,16,24). While several studies have characterized the infection response of different amphibian species (21,28–35), our work contributes to identify *Bd*’s functional machinery driving invasion and disease dynamics in a wide amphibian host range. In this study, we provide a framework to integrate available transcriptome data to further explore pathogen functional genomics during host-pathogen interactions. We expanded our view of a multi-host pathogen by characterizing the transcriptome landscapes of *Bd* across a diverse group of amphibians and inferred the origin and evolutionary history of pathogenic genes. Our results not only uncovered a conserved basal genetic machinery, but also highlight the functional capacity of *Bd* to display phenotypic plasticity in optimal and suboptimal host environments, which is decoupled from host phylogenetic relationships. Taken together, our findings indicate that gene expression plasticity is one of the evolutionary keys to the emergence of this panzootic multi-host pathogen.

Consistent with previous studies (22), *Bd* genetic machinery in host environments significantly differed from culture. Culture media is often a tryptone-based broth that provides *ad libitum* nutrients readily available to *Bd* for completing its lifecycle (16). This favorable condition leads to the expression of a high number of genes involved in numerous different biological processes. Likewise, our results show that *Bd* can easily acquire nutrients from the skin of susceptible anurans and urodeles for growing, developing, and, accordingly, express a large number of genes (Fig. 2). *Bd* expressed more sporangia-related genes in susceptible amphibians, indicating a more complex maturation, potentially reaching deeper skin layers. In these optimal environments, *Bd* could form complex vegetative and reproductive structures, such as densely branched rhizoids and clusters of thalli, which might contribute to disease progression and virulence (15). In contrast, in less susceptible hosts, *Bd* faces what we consider ‘challenging suboptimal environments’, which inflict nutritional restrictions related to limitations in substrate degradation, resulting in a restrained developmental capacity. In this scenario, *Bd* expressed a lower number of genes, perhaps involved in quick reproduction and protection, maybe through a resistant sporangium delaying the release of new zoospores, which has been suggested as a mechanism allowing the persistence and transmission of this emerging pathogen (25). Unlike the sister species *Bsal* that can form environmentally resistant non-motile spores (38), no resistant forms have been identified for *Bd* thus far. Variation in *Bd* life-history patterns has been demonstrated in culture under different thermal conditions (16,39). Likewise, endobiotic (inside the cell) and epibiotic (upon the cell surface) lifecycle strategies have been described *in vitro* observing the latest process in the explanted skin of a tolerant species (16). Additional evidence supporting plasticity is shown by the capacity of a closely related chytrid fungus to reallocate resources by altering rhizoid morphogenesis across environments with different resource availability (40). Our findings provide evidence for *Bd* gene expression plasticity, which could be underpin differences in lifecycles with variation in the mode and tempo of *Bd* development.

Gene expression plasticity can be adaptive or non-adaptive with evolutionary implications in both cases. Without empirical information about *Bd* fitness in different host environments, the identified conserved expression profile on each host environment can be a suitable indicator of adaptive plasticity. If the variation detected is adaptive, gene expression changes in response to host environments would enhance *Bd*’s ability to survive and reproduce in each host species. In addition, we hypothesized that similar biological traits arising from shared ancestry (i.e., immune defenses, behaviors, etc.) would lead to akin environmental pressures. However, we found no phylogenetic signal in *Bd* gene expression pattern, indicating that each amphibian species represents a unique independent environment. One potential explanation is that most of *Bd* isolates used to perform the experimental infections did not evolve with each host in nature representing novel interactions, hence the lack of phylogenetic signal supports the idea that plasticity predates co-evolutionary processes. Host environments also reflected intraspecific changes in susceptibility as our results confirmed variation in *Bd* gene expression in *P. cinereus* and *L. verreauxii alpine*, arising from different environmental conditions during exposures (i.e., temperature and disease status in nature, respectively (29,32). Thus, our analyses revealed high adaptability of *Bd*GPL and demonstrated that differences in host susceptibility are associated with changes in the fungal gene expression. Ultimately, these findings reflect gene expression reaction norms through trade-offs between virulence and transmission promoted by different host environments.

Our analyses expand previous knowledge about functional genomic elements for host exploitation and genomic innovations related to pathogenicity (19–22). Many of these genes encode extracellular carbohydrate-active enzymes part of the *Bd* secretome involved in biomass degradation (41), and were expressed across amphibians regardless of their susceptibility. When the nutritional substrate differed as tested in our *in vitro* versus *in vivo* comparisons, we expect a switch in enzyme composition, which as demonstrated by the increased expression of peptidases across all the studied amphibians compared to the tryptone-based broth (Fig. 2D and 2E). As other osmotrophic fungus after degrading the substrate, *Bd* needs to uptake nutrients, which requires membrane transports and transmembrane channels (16). We identified increased expression as well as signatures of positive selection in genes encoding transporters, which may be involved in scavenging nutrients and essential elements. Metals are one example of essential elements that provide virulence traits in many pathogenic fungi (e.g., cofactors of several peptidases) (42). They are also indispensable microelements for the amphibian hosts. Imbalance of metal homeostasis can be fatal for both pathogens and hosts, which compete in evolutionary arms race dynamics through pathogenesis and nutritional immunity, respectively (43). On the host side, nutritional immunity can involve diametrically opposite strategies: metal sequestration or toxicity via augmentation, or via metal-catalyzed generation of oxygen radicals. Pathogens must combat these strategies, for instance, by evolving efficient elimination, storage, and/or detoxification processes. The increased expression of genes encoding transporters, permeases, ion channels, and ubiquitination/deubiquitination enzymes provided evidence for the regulation of *Bd* homeostasis. Accordingly, competition for essential elements and changes in enzyme composition during the colonization of host environments could lead to infection progression and to the disruption of host homeostasis (44).

Genomic elements for host exploitation described hitherto were part of the common pathogenic machinery used by *Bd* to persist in a gradient of host environments (Fig. 1B). Our study provides an important contribution by characterizing *Bd* responses and to inferring the molecular evolution of pathogenic genes. In advantageous host environments, *Bd* increased the expression of not only additional peptidases supporting the capacity of *Bd* to better degrade these substrates, but also more adhesion and binding elements. These enzymes and elements confer the ability to attach, for instance, to leukocytes, which are known to be involved in the evasion of immune defenses (16). Indeed, increased pathogen virulence usually involves several mechanisms to sabotage host defenses (45). During the infection of susceptible hosts, *Bd* highly expressed regulatory proteins members of the Velvet complex involved in the production of secondary metabolites, which are bioactive molecules potentially harmful to the host (46). In contrast, the interactions with less susceptible hosts revealed a complex trans-kingdom communication with both host and pathogen interfering on each other, presumably through gene silencing (47). In these challenging scenarios, *Bd* could protect itself through dormancy of the zoospores or resistant sporangia. In addition, we found that *Bd* genetic armory differed in evolutionary origin, where the majority of the pathogenic genes were gained *de novo*, through expansion or horizontal gene transfer (HGT) events. In response to suboptimal host environments, most of the identified upregulated genes were candidates of HGT. These results are crucial to understand species interactions during the colonization process in the skin, where, for instance, host-associated microbes can alter infection outcome (48). As a strategy to survive, *Bd* could have stolen some of the genetic machinery from these microbes to combat resident bacteria with antifungal traits. In consequence, our findings linked how host environments present novel challenges and exert contrasting selective pressures in *Bd*. These results support the contribution of gene expression plasticity and the evolutionary innovations in the development of a very successful multi-host pathogen.

While some of the gene expression variation described in our study could be related to features of experimental design (e.g, *Bd* isolates, passage number (49), infection load), sequencing methodology (e.g, laser-capture microdissection or bulk RNA-Seq, sequencing coverage), stochastic changes associated with tissue harvest (e.g, cell location), and transcriptomic dynamics, our results unraveled plasticity arising via the interplay with different amphibian hosts. As a result, our findings provided an explanation of the host shifting ability of this pathogen. Additionally, our study yielded a prioritized list of genes to further investigate *Bd* responses in diverse and selective host environments. These genes could be used to design a microarray-based gene expression profiling to efficiently describe *Bd* plasticity and infection strategies across a much wider selection of amphibian hosts, including Asian species that have been persisting with the two chytrid fungi for a longer period of time (12). Future directions should include experiments to study *Bd* fitness in the different host environments, enabling empirical tests of virulence and transmission trade-offs. Likewise, we recommend functional analysis in parallel using dual RNA-Seq strategies (50) to have better estimates of how species interact with *Bd*, which can have important implications for amphibian conservation. Ultimately, our framework to analyze functional plasticity can help better understand the emergence and evolution of other multi-host pathogens.

## Material and Methods

### Infected amphibian skin and in culture transcriptome data

We selected RNA-Seq data from *Bd* in culture studies (21,22,31,33) and amphibian infection trials (21,22,28–30,32,34–37) and generated new infection transcriptome data for two amphibian species (see Table S1 for NCBI Short Read Archive accession codes and further information about transcriptome files). Data selection was made based on sequencing strategy (Illumina sequencing technology), experiment day, sequencing coverage, and number of biological replicates. Likewise, new transcriptomes from eleven skin samples of experimentally infected *Eleutherodactylus coqui* (36) and *Desmognathus auriculatus* (37) individuals were sequenced on an Illumina NovaSeq 6000 S4 2×150 flow cell after RNA extraction and cDNA library preparation using, respectively, QIAGEN RNeasy® Plus Mini Kit and NEBNext® Ultra™ II Directional RNA Library Prep Kit following manufacturers’ protocols. We measured the quantity and quality of the RNA and cDNA libraries using Qubit and Bioanalyzer. In total, we collated 145 transcriptomes encompassing samples from 14 different species. We classified the amphibian species by their susceptibility to *Bd* in three categories based on current literature (21,22,28–30,32,34–37): susceptible, partially susceptible, and tolerant hosts.

### Bd gene expression pattern analyses

Following the dual RNA-Seq approach (50), we recovered *Bd* expression profiles by in silico separating sequencing reads that mapped against the *Bd* JAM81 representative genome (GCF_000203795.1). From the newly generated transcriptomes, we previously filtered sequencing adaptors and trimmed the 10 left-bases of the reads using Trim Galore 0.6.5 (https://github.com/FelixKrueger/TrimGalore) and Prinseq 0.20.4 (51), respectively. The quality of the reads of all samples was checked using FastQC 0.11.5 (https://github.com/s-andrews/FastQC) after converting publicly available transcriptomes to fastq using SRA Toolkit fastq-dump (https://github.com/ncbi/sra-tools). We used gffread 0.11.8 (https://github.com/gpertea/gffread) to obtain the *Bd* genome gtf file and Star 2.7.3a (52) to count number reads per gene while mapping each file against the *Bd* reference genome (File S1 for raw count data). We normalized count data to account for sequencing depth and RNA composition using the median of ratios method of the R package DESeq2 (53) and explored sample similarity using a Principal Component Analysis (PCA). We visualized biological and technical sources of variation among samples by plotting the first five PCs. We tested the repeatability of the general *Bd* gene expression pattern across species using PC1 sample loadings with the R package rptR (54). To analyze *Bd* expression pattern in an amphibian phylogenetical framework, we retrieved 10,000 trees from VertLife (55) including the 14 host species, built a strict consensus tree, plotted it with FigTree v1.4.4 (https://github.com/rambaut/figtree/releases), and quantified phylogenetic signal by computing both Blomberg’s κ and Pagel’s λ for the species mean PC1 loadings using the R package phytools (56). We performed an ANOVA with Tukey’s honestly significant difference post hoc test to explore correlations between the species mean PC1 loadings and host susceptibility category.

### Differential gene expression analyses across a gradient of host susceptibility

To determine whether mean expression per gene of the different samples in each of the three host susceptibility categories was different from the mean of the *Bd* in culture samples, we fitted the count data to negative binomial models and performed Wald tests using the R package DESeq2 (53). We visualized common and unique differential expressed genes for each category using Euler diagrams with the R package eulerr (57). We annotated the upregulated genes (adjust p-value <= 0.05 and one or more than one positive unit of logarithmic fold expression change; log2FoldChange >= 1) against Uniprot (58) and Pfam (59) databases using BLAST 2.9.0 (60) and Hmmer 3.2.1 (61), respectively, and recovered GO and protein class information using retrieve/ID mapping of the Uniprot and Panther webtool (62), respectively. Additionally, we classified the upregulated genes in *Bd* sporangia or zoospore genes by following previously published differential gene expression analyses (31) between sporangia samples (SRR10389434, SRR10389435, SRR10389436) and zoospore samples (SRR10389437, SRR10389438, SRR10389439). To investigate the association between life-stage related genes and host susceptibility category, we compared the number of upregulated genes per susceptibility category that were annotated as sporangia genes using Pearson’s chi-square test and calculated Cramér’s V using the R package rcompanion (63).

### Molecular evolution analyses of chytrid genes

To infer the molecular evolution of the *Bd* upregulated genes on amphibian hosts, we analyzed the coding sequences (CDS) in a phylogenetic framework of 13 early-diverging zoosporic fungi species (see Table S3). After CDS translation of the genes of each genome using gffread 0.11.8 (https://github.com/gpertea/gffread), we inferred gene orthologs from aminoacidic sequences using OrthoFinder 2.3.11 (64), which allowed the identification of gene duplications and gene family expansion and contraction across species (number of genes in terminal branches greater than in internal ones and vice versa, respectively). After homolog discovery, we aligned the nucleotide sequences of the candidate orthogroups with more than 10 sequences that included any *Bd* upregulated genes using PRANK v.170427 (65). To detect genomic signatures of positive selection and evolutionary constraints of the *Bd* upregulated genes on amphibian hosts, we analyzed alignments and corresponding OrthoFinder gene trees using CODEML from PAML 4.8 (66) and conducted pairwise tests of nucleotide substitution models (Free Ratio vs M0 and MA vs nullMA) with their corresponding likelihood ratio tests (LRT).

### Data availability

The newly generated transcriptomic data have been deposited in the Sequence Read Archive, (BioProject accession number PRJNA739374). Previously published data were also used for this work and SRA codes for all studied samples can be found in Table S1. Complete code for the analyses including parameter specifications can be found in the File S2.

## Acknowledgments

We thank HiPerGator at the University of Florida and the National Center for Genome Analysis Support (NCGAS) at Indiana University for technical support and computational resources. We are also grateful to Iván de la Hera, Paula Silvar, George Glen, and Ariane Standing for valuable feedback. This work was supported by NSF Grant IOS-2011278 (to A.V.L), UF CLAS Research Office, and the Department of Biology startup funds. New samples contributed to this study were previously collected under NSF Dissertation Research Grant (IOS-1310036) and Postdoctoral Fellowship in Biology (DBI-1523551) to A.V.L. Undergraduate student J.V. was supported by the SF2UF Bridges Program (NIH-5R25GM115298-5).

## Authors Contributions

M.T.-S. and A.V.L. designed the research; M.T.-S., J.V., S. M.-B., and A.V.L. performed RNA extractions; M.T.-S. prepared cDNA libraries, analyzed the data, and wrote the original draft; all authors reviewed and approved the final manuscript.

## Supporting information

**Table S1**. Sample information. Transcriptome data information from where we recovered the 145 *Batrachochytrium dendrobatidis* (*Bd*) gene expression profiles. Samples obtained from in culture have no host genera, order, and susceptibility category information (-).

**Table S2**. The explained proportion of the total variance by each principal component. We assessed sample similarity using Principal Component Analysis (PCA) and identified PC clustering gene expression profiles.

**Table S3**. Fungal genome information. For each of the 13 early-divergent zoosporic fungi, taxonomy information from the NCBI taxonomy browser and NCBI accession numbers are provided.

**Figure S1**. Principal Component Analysis of *Bd* gene expression profiles for 145 samples of infected skin and cultures. Object size illustrates sequencing coverage (Megabases, Mb) and color refers to sequencing project while object shapes differentiate the origin of the samples: amphibian skin (•) and in culture (▴).

**Figure S2**. Principal Component Analysis of *Bd* gene expression profiles for 145 samples of infected skin and cultures. Object color illustrates *Bd* strain used to experimentally infect amphibians of the different studies while object shapes differentiate the origin of the samples: amphibian skin (•) and in culture (▴).

**Figure S3**. Mean PC1 loadings per amphibian species and host susceptibility category. The boxplot displays pairwise comparisons of Bd gene expression pattern per susceptibility category. Asterisks denote significant comparisons.

**Figure S4**. Phylogenetic tree of 13 early-divergent zoosporic fungi showing duplications for the identified gene families. Branches are color-coded based in terminal gene duplications.

**Figure S5**. Numeric distribution of the nucleotide substitutions ratios (dN/dS) of the gene families including Bd pathogenic genes. Density plots are color-coded by type of candidate gene family symbolizing the number of gene families by bars on the x-axis.

**File S1 (separate file)**. *Batrachochytrium dendrobatidis* (*Bd*) gene expression information. Excel file with two sheets containing raw read counts for the 145 samples (sheet 1: raw_counts) and upregulated genes with their annotations and gene family information (sheet 2: upregulated_genes). Raw counts are displayed in a table manner with *Bd* counts per gene (8700 genes included in the *Bd* JAM81 representative genome, GCF_000203795.1) in each row for the 145 samples with their SRA codes in each column. For the upregulated genes, each gene annotations include NCBI accession numbers (NCBI_accession_numbers), Uniport best hit information (Uniprot_id and Uniprot_protein_name), Pfam best hit information (Pfam_id and Pfam_description), and Gene Ontology domains description (GO_biological_process, GO_cellular_component, and GO_molecular_function). In this second excel sheet, we included information from the differential expression analyses: mean expression across samples (BaseMean_expression); fold change in logarithmic scale for the contrasts susceptible host versus in culture (log2FC_Susceptible), partially susceptible host versus in culture (log2FC_Partial_Susceptible), and tolerant hosts versus in culture (log2FC_Tolerant); classification from the contrast zoospore versus sporangia (Life_stage); and gene pathogenic category based on the decision tree represented in Fig. 1B (Pathogenic_category). Additionally, we reported information of the gene family (Orthogroup_id) including the number of both *Bd* and total genes in each gene family (Number_Bd_genes and Total_gene_number, respectively) and molecular evolution: categorization based on origin and evolutionary history (Gene_familie_category), nucleotide substitution ratio under M0 and MA evolutionary models (M0_dN/dS and MA_2a_foreground_dN/dS, respectively).

**File S2 (separate file)**. Complete code for all analyses presented in the article.

