## Supplementary material for "Panzootic chytrid fungus exploits diverse amphibian host environments through plastic infection strategies": Tables S1 to S3; Figures S1 to S5

**This PDF file includes:**

Tables S1 to S3

Figures S1 to S5

Legends for File S1 to S2

**Other supplementary materials for this manuscript include the following:**

File S1

File S2

**Table S1**. Sample information. Transcriptome data information from where we recovered the 145 *Batrachochytrium dendrobatidis* (*Bd*) gene expression profiles. Samples obtained from in culture have no host genera, order, and susceptibility category information (-).

| SRA file | BioProject | Species | Host genera | Host order | Host susceptibility category | *Bd* isolate | *Bd* genes |
|---|---|---|---|---|---|---|---|
| SRR10389434 | PRJNA563776 | *Batrachochytrium dendrobatidis* | - | - | - | VM1 | 7780 |
| SRR10389435 | PRJNA563776 | *Batrachochytrium dendrobatidis* | - | - | - | VM1 | 7687 |
| SRR10389436 | PRJNA563776 | *Batrachochytrium dendrobatidis* | - | - | - | VM1 | 7697 |
| SRR10389437 | PRJNA563776 | *Batrachochytrium dendrobatidis* | - | - | - | VM1 | 7446 |
| SRR10389438 | PRJNA563776 | *Batrachochytrium dendrobatidis* | - | - | - | VM1 | 7430 |
| SRR10389439 | PRJNA563776 | *Batrachochytrium dendrobatidis* | - | - | - | VM1 | 7536 |
| SRR10389440 | PRJNA563776 | *Batrachochytrium dendrobatidis* | - | - | - | VM1 | 7773 |
| SRR10389441 | PRJNA563776 | *Batrachochytrium dendrobatidis* | - | - | - | VM1 | 7774 |
| SRR10389442 | PRJNA563776 | *Batrachochytrium dendrobatidis* | - | - | - | VM1 | 7770 |
| SRR11309525 | PRJNA612733 | *Notophthalmus viridescens* | *Notophthalmus* | Urodela | Tolerant | Natural | 710 |
| SRR11309526 | PRJNA612733 | *Notophthalmus viridescens* | *Notophthalmus* | Urodela | Tolerant | Natural | 486 |
| SRR11309527 | PRJNA612733 | *Notophthalmus viridescens* | *Notophthalmus* | Urodela | Tolerant | Natural | 269 |
| SRR11309536 | PRJNA612733 | *Notophthalmus viridescens* | *Notophthalmus* | Urodela | Tolerant | Natural | 281 |
| SRR11880849 | PRJNA636076 | *Lithobates yavapaiensis* | *Lithobates* | Anura | Partially susceptible | JEL423 | 1 |
| SRR11880859 | PRJNA636076 | *Lithobates yavapaiensis* | *Lithobates* | Anura | Partially susceptible | JEL423 | 42 |
| SRR11880867 | PRJNA636076 | *Lithobates yavapaiensis* | *Lithobates* | Anura | Partially susceptible | JEL423 | 2 |
| SRR11880877 | PRJNA636076 | *Lithobates yavapaiensis* | *Lithobates* | Anura | Partially susceptible | JEL423 | 1 |
| SRR11880903 | PRJNA636076 | *Lithobates yavapaiensis* | *Lithobates* | Anura | Partially susceptible | JEL423 | 1 |
| SRR11880909 | PRJNA636076 | *Lithobates yavapaiensis* | *Lithobates* | Anura | Partially susceptible | JEL423 | 2 |
| SRR11880929 | PRJNA636076 | *Lithobates yavapaiensis* | *Lithobates* | Anura | Partially susceptible | JEL423 | 104 |
| SRR11880936 | PRJNA636076 | *Lithobates yavapaiensis* | *Lithobates* | Anura | Partially susceptible | JEL423 | 4 |
| SRR11880941 | PRJNA636076 | *Lithobates yavapaiensis* | *Lithobates* | Anura | Partially susceptible | JEL423 | 114 |
| SRR11880951 | PRJNA636076 | *Lithobates yavapaiensis* | *Lithobates* | Anura | Partially susceptible | JEL423 | 2 |
| SRR11880956 | PRJNA636076 | *Lithobates yavapaiensis* | *Lithobates* | Anura | Partially susceptible | JEL423 | 2 |
| SRR1560902 | PRJNA259742 | *Craugastor fitzingeri* | *Craugastor* | Anura | Tolerant | JEL423 | 522 |
| SRR1560903 | PRJNA259742 | *Craugastor fitzingeri* | *Craugastor* | Anura | Tolerant | JEL423 | 1956 |
| SRR1560906 | PRJNA259742 | *Craugastor fitzingeri* | *Craugastor* | Anura | Tolerant | JEL423 | 2798 |
| SRR1560909 | PRJNA259742 | *Craugastor fitzingeri* | *Craugastor* | Anura | Tolerant | JEL423 | 3449 |
| SRR1560910 | PRJNA259742 | *Craugastor fitzingeri* | *Craugastor* | Anura | Tolerant | JEL423 | 4699 |
| SRR1560915 | PRJNA259742 | *Craugastor fitzingeri* | *Craugastor* | Anura | Tolerant | JEL423 | 137 |
| SRR1560992 | PRJNA259743 | *Agalychnis callidryas* | *Agalychnis* | Anura | Tolerant | JEL423 | 92 |
| SRR1560994 | PRJNA259743 | *Agalychnis callidryas* | *Agalychnis* | Anura | Tolerant | JEL423 | 65 |
| SRR1560999 | PRJNA259743 | *Agalychnis callidryas* | *Agalychnis* | Anura | Tolerant | JEL423 | 226 |
| SRR1561000 | PRJNA259743 | *Agalychnis callidryas* | *Agalychnis* | Anura | Tolerant | JEL423 | 67 |
| SRR1562821 | PRJNA259720 | *Atelopus glyphus* | *Atelopus* | Anura | Susceptible | JEL423 | 6997 |
| SRR1562822 | PRJNA259720 | *Atelopus glyphus* | *Atelopus* | Anura | Susceptible | JEL423 | 7016 |
| SRR1562823 | PRJNA259720 | *Atelopus glyphus* | *Atelopus* | Anura | Susceptible | JEL423 | 7366 |
| SRR1562824 | PRJNA259720 | *Atelopus glyphus* | *Atelopus* | Anura | Susceptible | JEL423 | 7033 |
| SRR1562825 | PRJNA259720 | *Atelopus glyphus* | *Atelopus* | Anura | Susceptible | JEL423 | 3135 |
| SRR1562831 | PRJNA259720 | *Atelopus glyphus* | *Atelopus* | Anura | Susceptible | JEL423 | 6208 |

| | | | | | | | |
|---|---|---|---|---|---|---|---|
| SRR1562832 | PRJNA259720 | *Atelopus glyphus* | *Atelopus* | Anura | Susceptible | JEL423 | 5211 |
| SRR1562835 | PRJNA259720 | *Atelopus glyphus* | *Atelopus* | Anura | Susceptible | JEL423 | 6314 |
| SRR2719455 | PRJNA299101 | *Batrachochytrium dendrobatidis* | - | - | - | JEL423 | 7881 |
| SRR2719581 | PRJNA299101 | *Batrachochytrium dendrobatidis* | - | - | - | JEL423 | 7851 |
| SRR2719582 | PRJNA299101 | *Batrachochytrium dendrobatidis* | - | - | - | JEL423 | 7883 |
| SRR2719583 | PRJNA299101 | *Batrachochytrium dendrobatidis* | - | - | - | JEL423 | 7896 |
| SRR2720403 | PRJNA299101 | *Atelopus zeteki* | *Atelopus* | Anura | Susceptible | JEL423 | 7487 |
| SRR2720404 | PRJNA299101 | *Atelopus zeteki* | *Atelopus* | Anura | Susceptible | JEL423 | 7996 |
| SRR2720406 | PRJNA299101 | *Atelopus zeteki* | *Atelopus* | Anura | Susceptible | JEL423 | 7935 |
| SRR2720407 | PRJNA299101 | *Atelopus zeteki* | *Atelopus* | Anura | Susceptible | JEL423 | 7795 |
| SRR2721696 | PRJNA299101 | *Agalychnis lemur* | *Agalychnis* | Anura | Susceptible | JEL423 | 7474 |
| SRR2721749 | PRJNA299101 | *Agalychnis lemur* | *Agalychnis* | Anura | Susceptible | JEL423 | 7664 |
| SRR2723982 | PRJNA299101 | *Agalychnis lemur* | *Agalychnis* | Anura | Susceptible | JEL423 | 2686 |
| SRR2724026 | PRJNA299101 | *Agalychnis lemur* | *Agalychnis* | Anura | Susceptible | JEL423 | 6874 |
| SRR3048025 | PRJNA300849 | *Tylototriton wenxianensis* | *Tylototriton* | Urodela | Susceptible | JEL423 | 2820 |
| SRR3048769 | PRJNA300849 | *Tylototriton wenxianensis* | *Tylototriton* | Urodela | Susceptible | JEL423 | 6316 |
| SRR3048772 | PRJNA300849 | *Tylototriton wenxianensis* | *Tylototriton* | Urodela | Susceptible | JEL423 | 5491 |
| SRR3707435 | PRJNA326253 | *Batrachochytrium dendrobatidis* | - | - | - | JEL423 | 8048 |
| SRR3707957 | PRJNA326253 | *Batrachochytrium dendrobatidis* | - | - | - | JEL423 | 8236 |
| SRR5149382 | PRJNA356986 | *Litoria verreauxii alpina* | *Litoria* | Anura | Partially susceptible | AbercrombieNP-L.booroolongensis-09-LB-P7 | 118 |
| SRR5149384 | PRJNA356986 | *Litoria verreauxii alpina* | *Litoria* | Anura | Partially susceptible | AbercrombieNP-L.booroolongensis-09-LB-P7 | 7213 |
| SRR5149397 | PRJNA356986 | *Litoria verreauxii alpina* | *Litoria* | Anura | Partially susceptible | AbercrombieNP-L.booroolongensis-09-LB-P7 | 2384 |
| SRR5149398 | PRJNA356986 | *Litoria verreauxii alpina* | *Litoria* | Anura | Partially susceptible | AbercrombieNP-L.booroolongensis-09-LB-P7 | 6519 |
| SRR5149441 | PRJNA356986 | *Litoria verreauxii alpina* | *Litoria* | Anura | Partially susceptible | AbercrombieNP-L.booroolongensis-09-LB-P7 | 477 |
| SRR5149450 | PRJNA356986 | *Litoria verreauxii alpina* | *Litoria* | Anura | Partially susceptible | AbercrombieNP-L.booroolongensis-09-LB-P7 | 81 |
| SRR5149451 | PRJNA356986 | *Litoria verreauxii alpina* | *Litoria* | Anura | Partially susceptible | AbercrombieNP-L.booroolongensis-09-LB-P7 | 4240 |
| SRR5149493 | PRJNA356986 | *Litoria verreauxii alpina* | *Litoria* | Anura | Partially susceptible | AbercrombieNP-L.booroolongensis-09-LB-P7 | 897 |
| SRR5149496 | PRJNA356986 | *Litoria verreauxii alpina* | *Litoria* | Anura | Partially susceptible | AbercrombieNP-L.booroolongensis-09-LB-P7 | 1411 |
| SRR5149518 | PRJNA356986 | *Litoria verreauxii alpina* | *Litoria* | Anura | Partially susceptible | AbercrombieNP-L.booroolongensis-09-LB-P7 | 5421 |
| SRR5149535 | PRJNA356986 | *Litoria verreauxii alpina* | *Litoria* | Anura | Partially susceptible | AbercrombieNP-L.booroolongensis-09-LB-P7 | 148 |
| SRR5149542 | PRJNA356986 | *Litoria verreauxii alpina* | *Litoria* | Anura | Partially susceptible | AbercrombieNP-L.booroolongensis-09-LB-P7 | 6634 |

| | | | | | | | |
|---|---|---|---|---|---|---|---|
| SRR5810415 | PRJNA392411 | *Lithobates catesbeianus* | *Lithobates* | Anura | Tolerant | Section Line | 292 |
| SRR5810416 | PRJNA392411 | *Lithobates catesbeianus* | *Lithobates* | Anura | Tolerant | Section Line | 30 |
| SRR5810417 | PRJNA392411 | *Lithobates catesbeianus* | *Lithobates* | Anura | Tolerant | Carter Meadow | 65 |
| SRR5810429 | PRJNA392411 | *Lithobates catesbeianus* | *Lithobates* | Anura | Tolerant | Carter Meadow | 227 |
| SRR5810436 | PRJNA392411 | *Lithobates catesbeianus* | *Lithobates* | Anura | Tolerant | Section Line | 31 |
| SRR5810437 | PRJNA392411 | *Lithobates catesbeianus* | *Lithobates* | Anura | Tolerant | Section Line | 116 |
| SRR5810439 | PRJNA392411 | *Lithobates sylvatica* | *Lithobates* | Anura | Partially susceptible | Section Line | 2731 |
| SRR5810441 | PRJNA392411 | *Lithobates sylvatica* | *Lithobates* | Anura | Partially susceptible | Carter Meadow | 18 |
| SRR5810444 | PRJNA392411 | *Lithobates sylvatica* | *Lithobates* | Anura | Partially susceptible | Section Line | 156 |
| SRR5810445 | PRJNA392411 | *Lithobates sylvatica* | *Lithobates* | Anura | Partially susceptible | Section Line | 2755 |
| SRR5810450 | PRJNA392411 | *Lithobates sylvatica* | *Lithobates* | Anura | Partially susceptible | Carter Meadow | 303 |
| SRR5810451 | PRJNA392411 | *Lithobates sylvatica* | *Lithobates* | Anura | Partially susceptible | Carter Meadow | 53 |
| SRR5810455 | PRJNA392411 | *Lithobates sylvatica* | *Lithobates* | Anura | Partially susceptible | Section Line | 3194 |
| SRR5810456 | PRJNA392411 | *Lithobates sylvatica* | *Lithobates* | Anura | Partially susceptible | Section Line | 297 |
| SRR5810467 | PRJNA392411 | *Lithobates sylvatica* | *Lithobates* | Anura | Partially susceptible | Carter Meadow | 500 |
| SRR5810476 | PRJNA392411 | *Lithobates sylvatica* | *Lithobates* | Anura | Partially susceptible | Section Line | 147 |
| SRR5810477 | PRJNA392411 | *Lithobates sylvatica* | *Lithobates* | Anura | Partially susceptible | Section Line | 1913 |
| SRR5810482 | PRJNA392411 | *Lithobates sylvatica* | *Lithobates* | Anura | Partially susceptible | Section Line | 780 |
| SRR5810483 | PRJNA392411 | *Lithobates sylvatica* | *Lithobates* | Anura | Partially susceptible | Section Line | 158 |
| SRR5810485 | PRJNA392411 | *Lithobates sylvatica* | *Lithobates* | Anura | Partially susceptible | Carter Meadow | 325 |
| SRR5810491 | PRJNA392411 | *Lithobates catesbeianus* | *Lithobates* | Anura | Tolerant | Carter Meadow | 31 |
| SRR5810494 | PRJNA392411 | *Lithobates catesbeianus* | *Lithobates* | Anura | Tolerant | Section Line | 112 |
| SRR5810495 | PRJNA392411 | *Lithobates catesbeianus* | *Lithobates* | Anura | Tolerant | Carter Meadow | 34 |
| SRR5810496 | PRJNA392411 | *Lithobates catesbeianus* | *Lithobates* | Anura | Tolerant | Carter Meadow | 23 |
| SRR5988742 | PRJNA400613 | *Batrachochytrium dendrobatidis* | - | - | - | CLFT044 | 8083 |
| SRR5988743 | PRJNA400613 | *Batrachochytrium dendrobatidis* | - | - | - | CLFT044 | 8121 |
| SRR5988744 | PRJNA400613 | *Batrachochytrium dendrobatidis* | - | - | - | JEL410 | 7878 |
| SRR5988745 | PRJNA400613 | *Batrachochytrium dendrobatidis* | - | - | - | JEL410 | 7908 |
| SRR5988746 | PRJNA400613 | *Batrachochytrium dendrobatidis* | - | - | - | CLFT001 | 7995 |
| SRR5988747 | PRJNA400613 | *Batrachochytrium dendrobatidis* | - | - | - | CLFT001 | 8045 |
| SRR5988748 | PRJNA400613 | *Batrachochytrium dendrobatidis* | - | - | - | JEL422 | 7652 |
| SRR5988749 | PRJNA400613 | *Batrachochytrium dendrobatidis* | - | - | - | JEL422 | 7825 |
| SRR5988750 | PRJNA400613 | *Batrachochytrium dendrobatidis* | - | - | - | CLFT001 | 8104 |
| SRR5988751 | PRJNA400613 | *Batrachochytrium dendrobatidis* | - | - | - | CLFT023 | 7895 |
| SRR5988752 | PRJNA400613 | *Batrachochytrium dendrobatidis* | - | - | - | CLFT026 | 7909 |
| SRR5988753 | PRJNA400613 | *Batrachochytrium dendrobatidis* | - | - | - | CLFT044 | 8091 |
| SRR5988754 | PRJNA400613 | *Batrachochytrium dendrobatidis* | - | - | - | CLFT023 | 7991 |
| SRR5988755 | PRJNA400613 | *Batrachochytrium dendrobatidis* | - | - | - | CLFT026 | 7998 |
| SRR5988756 | PRJNA400613 | *Batrachochytrium dendrobatidis* | - | - | - | JEL422 | 7600 |
| SRR5988757 | PRJNA400613 | *Batrachochytrium dendrobatidis* | - | - | - | CLFT023 | 8007 |
| SRR5988758 | PRJNA400613 | *Batrachochytrium dendrobatidis* | - | - | - | CLFT026 | 7862 |
| SRR5988759 | PRJNA400613 | *Batrachochytrium dendrobatidis* | - | - | - | JEL410 | 7915 |
| SRR955792 | PRJNA216050 | *Atelopus zeteki* | *Atelopus* | Anura | Susceptible | JEL423 | 6383 |
| SRR955793 | PRJNA216050 | *Atelopus zeteki* | *Atelopus* | Anura | Susceptible | JEL423 | 6035 |
| SRR955794 | PRJNA216050 | *Atelopus zeteki* | *Atelopus* | Anura | Susceptible | JEL423 | 7318 |
| SRR955795 | PRJNA216050 | *Atelopus zeteki* | *Atelopus* | Anura | Susceptible | JEL423 | 7155 |
| SRR957174 | PRJNA216050 | *Atelopus zeteki* | *Atelopus* | Anura | Susceptible | JEL423 | 5441 |
| SRR957175 | PRJNA216050 | *Atelopus zeteki* | *Atelopus* | Anura | Susceptible | JEL423 | 6976 |

| | | | | | | | |
|---|---|---|---|---|---|---|---|
| SRR9925252 | PRJNA559247 | *Plethodon cinereus* | *Plethodon* | Urodela | Tolerant | JEL423 | 116 |
| SRR9925255 | PRJNA559247 | *Plethodon cinereus* | *Plethodon* | Urodela | Tolerant | JEL423 | 731 |
| SRR9925257 | PRJNA559247 | *Plethodon cinereus* | *Plethodon* | Urodela | Tolerant | JEL423 | 90 |
| SRR9925259 | PRJNA559247 | *Plethodon cinereus* | *Plethodon* | Urodela | Tolerant | JEL423 | 128 |
| SRR9925260 | PRJNA559247 | *Plethodon cinereus* | *Plethodon* | Urodela | Tolerant | JEL423 | 2308 |
| SRR9925261 | PRJNA559247 | *Plethodon cinereus* | *Plethodon* | Urodela | Tolerant | JEL423 | 574 |
| SRR9925277 | PRJNA559247 | *Plethodon cinereus* | *Plethodon* | Urodela | Tolerant | JEL423 | 4248 |
| SRR9925278 | PRJNA559247 | *Plethodon cinereus* | *Plethodon* | Urodela | Tolerant | JEL423 | 76 |
| SRR9925279 | PRJNA559247 | *Plethodon cinereus* | *Plethodon* | Urodela | Tolerant | JEL423 | 56 |
| SRR9925280 | PRJNA559247 | *Plethodon cinereus* | *Plethodon* | Urodela | Tolerant | JEL423 | 62 |
| SRR9925282 | PRJNA559247 | *Plethodon cinereus* | *Plethodon* | Urodela | Tolerant | JEL423 | 68 |
| SRR9925285 | PRJNA559247 | *Plethodon cinereus* | *Plethodon* | Urodela | Tolerant | JEL423 | 1305 |
| SRR9925288 | PRJNA559247 | *Plethodon cinereus* | *Plethodon* | Urodela | Tolerant | JEL423 | 5292 |
| SRR9925289 | PRJNA559247 | *Plethodon cinereus* | *Plethodon* | Urodela | Tolerant | JEL423 | 1135 |
| SRR9925300 | PRJNA559247 | *Plethodon cinereus* | *Plethodon* | Urodela | Tolerant | JEL423 | 546 |
| SRR14866077 | PRJNA739374 | *Eleutherodactylus coqui* | *Eleutherodactylus* | Anura | Partially susceptible | JEL427 | 28 |
| SRR14866076 | PRJNA739374 | *Eleutherodactylus coqui* | *Eleutherodactylus* | Anura | Partially susceptible | JEL427 | 150 |
| SRR14866074 | PRJNA739374 | *Eleutherodactylus coqui* | *Eleutherodactylus* | Anura | Partially susceptible | JEL427 | 52 |
| SRR14866073 | PRJNA739374 | *Eleutherodactylus coqui* | *Eleutherodactylus* | Anura | Partially susceptible | JEL427 | 42 |
| SRR14866072 | PRJNA739374 | *Eleutherodactylus coqui* | *Eleutherodactylus* | Anura | Partially susceptible | JEL427 | 113 |
| SRR14866071 | PRJNA739374 | *Eleutherodactylus coqui* | *Eleutherodactylus* | Anura | Partially susceptible | JEL427 | 62 |
| SRR14866070 | PRJNA739374 | *Eleutherodactylus coqui* | *Eleutherodactylus* | Anura | Partially susceptible | JEL427 | 140 |
| SRR14866069 | PRJNA739374 | *Eleutherodactylus coqui* | *Eleutherodactylus* | Anura | Partially susceptible | JEL427 | 26 |
| SRR14866068 | PRJNA739374 | *Eleutherodactylus coqui* | *Eleutherodactylus* | Anura | Partially susceptible | JEL427 | 29 |
| SRR14866067 | PRJNA739374 | *Eleutherodactylus coqui* | *Eleutherodactylus* | Anura | Partially susceptible | JEL427 | 20 |
| SRR14866075 | PRJNA739374 | *Desmognathus auriculatus* | *Desmognathus* | Urodela | Susceptible | ALKL1 | 3140 |

**Table S2**. The explained proportion of the total variance by each principal component. We assessed sample similarity using Principal Component Analysis (PCA) and identified PC clustering gene expression profiles.

| | PC1 | PC2 | PC3 | PC4 | PC5 | PC6 |
|---|---|---|---|---|---|---|
| Proportion of Variance | 70.87 | 5.15 | 2.46 | 1.92 | 1.48 | 0.94 |
| Cumulative Proportion | 70.87 | 76.03 | 78.49 | 80.41 | 81.89 | 82.86 |

**Table S3**. Fungal genome information. For each of the 13 early-divergent zoosporic fungi, taxonomy information from the NCBI taxonomy browser and NCBI accession numbers are provided.

| **Species** | **Order** | **Class** | **Division** | **Accession numbers** |
|---|---|---|---|---|
| *Batrachochytrium dendrobatidis* | Rhizophydiales | Chytridiomycetes | Chytridiomycota | GCF_000203795.1 |
| *Batrachochytrium salamandrivorans* | Rhizophydiales | Chytridiomycetes | Chytridiomycota | GCA_002006685.1 |
| *Blyttiomyces helicus* | Rhizophlyctidales | Chytridiomycetes | Chytridiomycota | GCA_003614705.1 |
| *Powellomyces hirtus* | Spizellomycetales | Chytridiomycetes | Chytridiomycota | GCA_006536005.1 |
| *Spizellomyces punctatus* | Spizellomycetales | Chytridiomycetes | Chytridiomycota | GCF_000182565.1 |
| *Rhizoclosmatium globosum* | Chytridiales | Chytridiomycetes | Chytridiomycota | GCA_002104985.1 |
| *Chytriomyces confervae* | Chytridiales | Chytridiomycetes | Chytridiomycota | GCA_006535975.1 |
| *Synchytrium microbalum* | Synchytriales | Chytridiomycetes | Chytridiomycota | GCF_006535985.1 |
| *Caulochytrium protostelioides* | Caulochytriales | Chytridiomycetes | Chytridiomycota | GCA_003615045.1 |
| *Gonapodya prolifera* | Monoblepharidales | Monoblepharidomycetes | Chytridiomycota | GCA_001574975.1 |
| *Anaeromyces robustus* | Neocallimastigales | Neocallimastigomycetes | Chytridiomycota | GCA_002104895.1 |
| *Neocallimastix californiae* | Neocallimastigales | Neocallimastigomycetes | Chytridiomycota | GCA_002104975.1 |
| *Piromyces finnis* | Neocallimastigales | Neocallimastigomycetes | Chytridiomycota | GCA_002104945.1 |

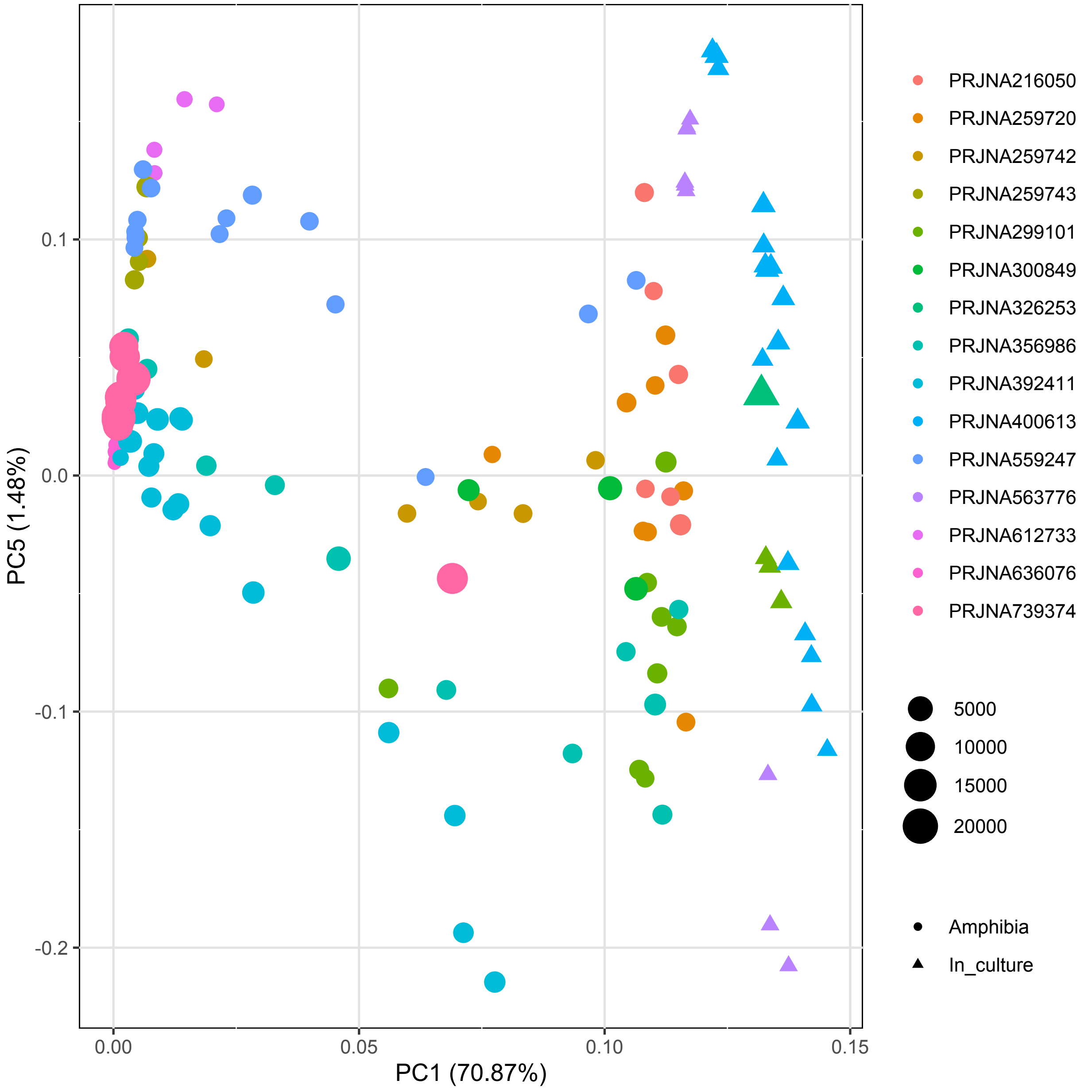


**Figure S1**. Principal Component Analysis of *Bd* gene expression profiles for 145 samples of infected skin and cultures. Object size illustrates sequencing coverage (Megabases, Mb) and color refers to sequencing project while object shapes differentiate the origin of the samples: amphibian skin (●) and in culture (▲).

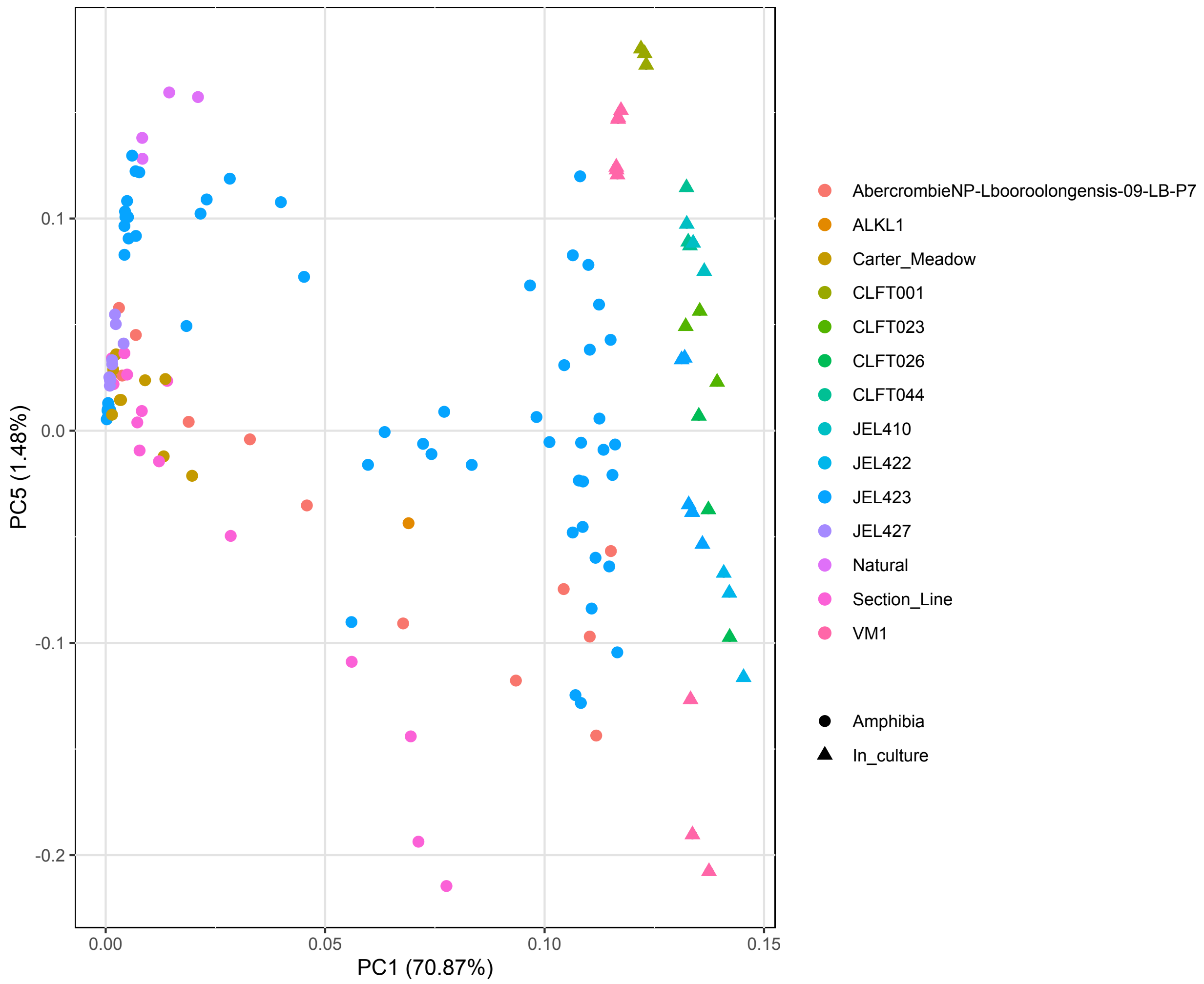


**Figure S2**. Principal Component Analysis of *Bd* gene expression profiles for 145 samples of infected skin and cultures. Object color illustrates *Bd* strain used to experimentally infect amphibians of the different studies while object shapes differentiate the origin of the samples: amphibian skin (●) and in culture (▲).

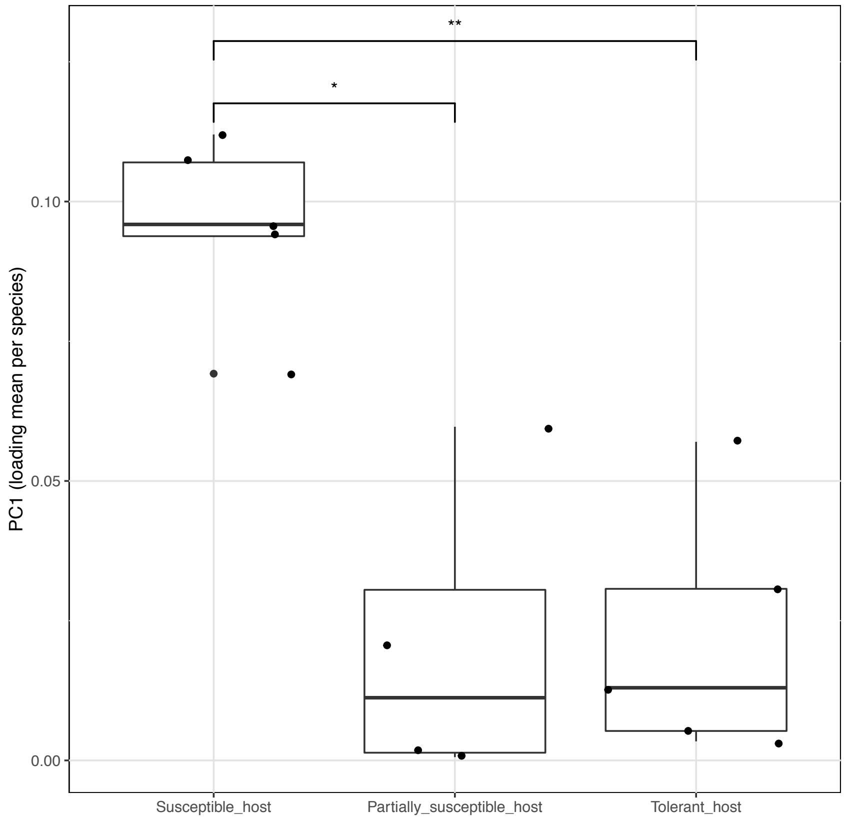


**Figure S3**. Mean PC1 loadings per amphibian species and host susceptibility category. The boxplot displays pairwise comparisons of Bd gene expression pattern per susceptibility category. Each dot represents one amphibian species. Asterisks denote significant comparisons of the Tukey's honestly significant difference post hoc test (alpha < 0.05).

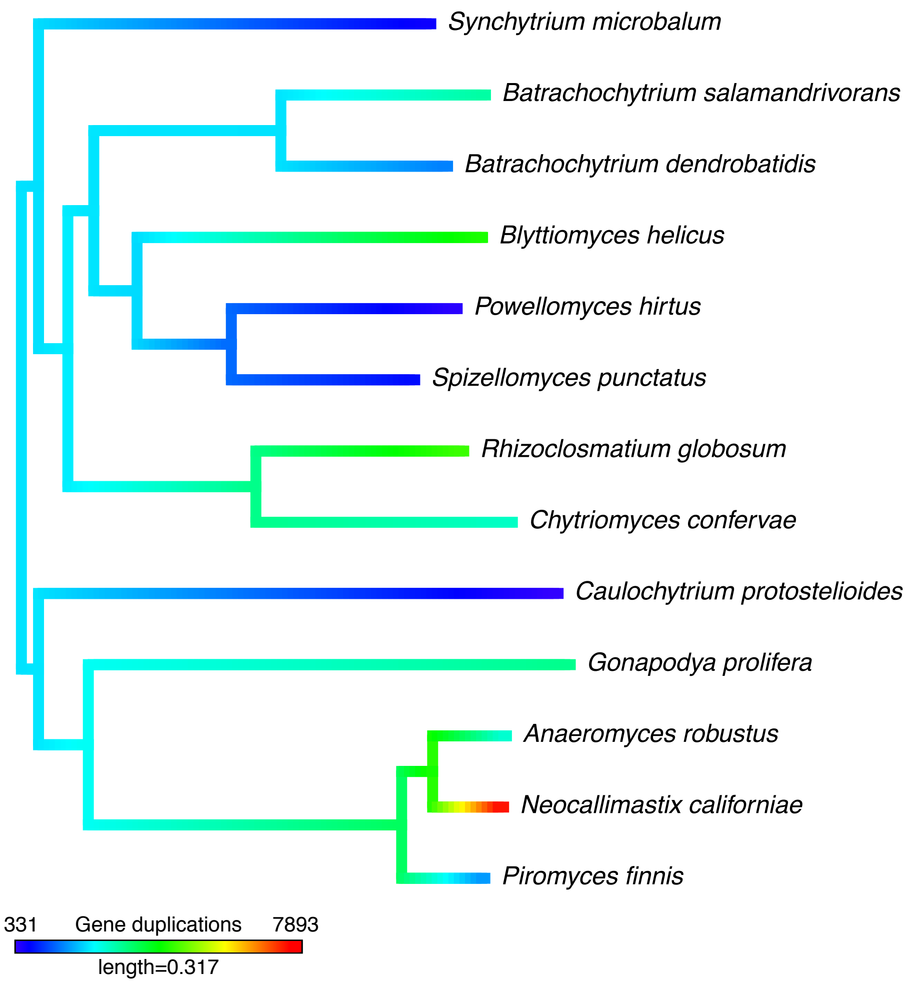


**Figure S4**. Phylogenetic tree of 13 early-divergent zoosporic fungi showing duplications for the identified gene families. Branches are color-coded based in terminal gene duplications.

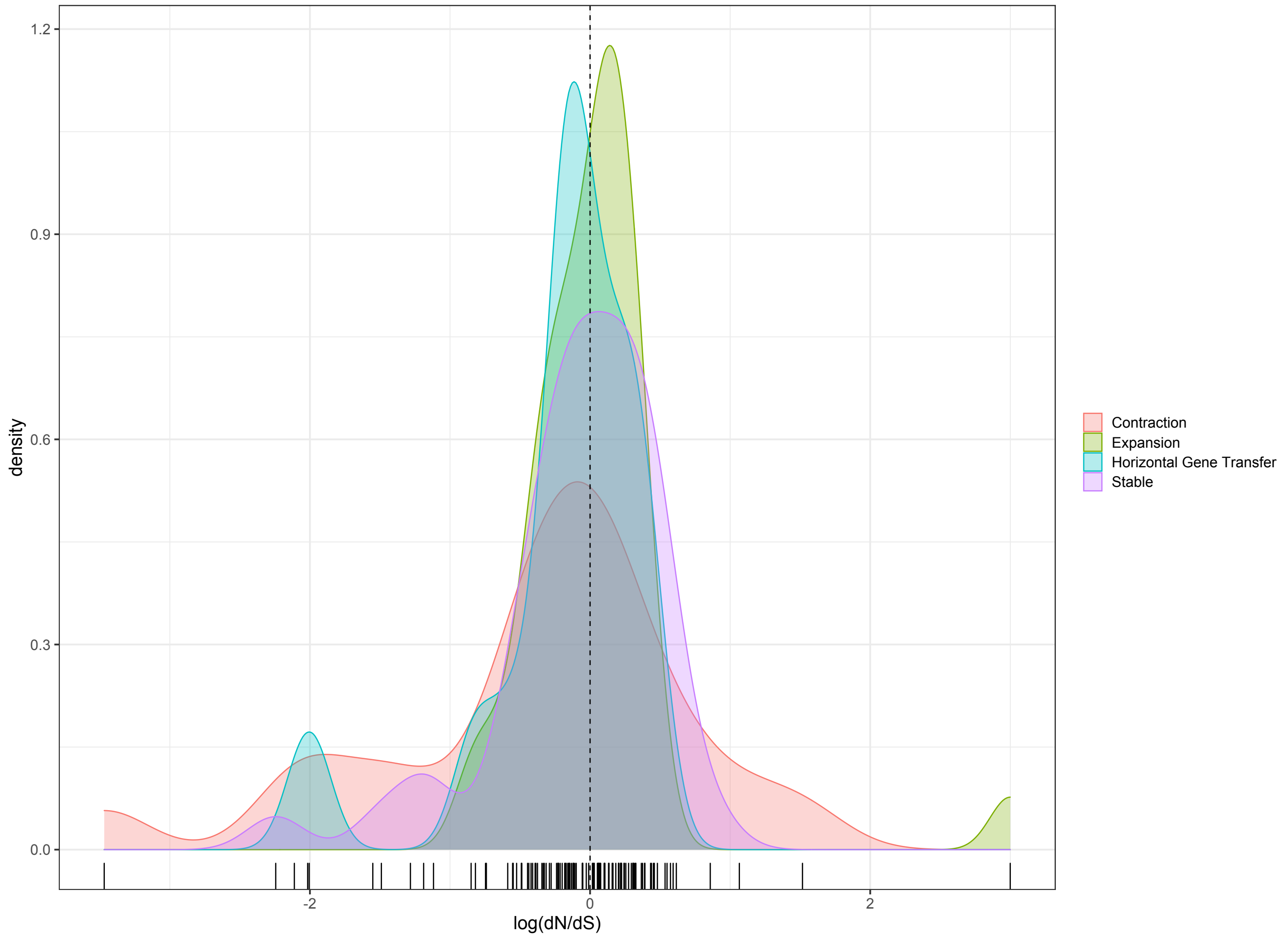


**Figure S5.** Distribution of nucleotide substitutions ratios (dN/dS) of the gene families including Bd pathogenic genes. Density plots are color-coded by type of candidate gene family symbolizing the number of gene families by bars on the x-axis.

**File S1 (separate file)**. *Batrachochytrium dendrobatidis* (*Bd*) gene expression information. Excel file with two sheets containing raw read counts for the 145 samples (sheet 1: raw_counts) and upregulated genes with their annotations and gene family information (sheet 2: upregulated_genes). Raw counts are displayed in a table manner with *Bd* counts per gene (8700 genes included in the *Bd* JAM81 representative genome, GCF_000203795.1) in each row for the 145 samples with their SRA codes in each column. For the upregulated genes, each gene annotations include NCBI accession numbers (NCBI_accession_numbers), Uniport best hit information (Uniprot_id and Uniprot_protein_name), Pfam best hit information (Pfam_id and Pfam_description), and Gene Ontology domains description (GO_biological_process, GO_cellular_component, and GO_molecular_function). In this second excel sheet, we included information from the differential expression analyses: mean expression across samples (BaseMean_expression); fold change in logarithmic scale for the contrasts susceptible host versus in culture (log2FC_Susceptible), partially susceptible host versus in culture (log2FC_Partial_Susceptible), and tolerant hosts versus in culture (log2FC_Tolerant); classification from the contrast zoospore versus sporangia (Life_stage); and gene pathogenic category based on the decision tree represented in Fig. 1B (Pathogenic_category). Additionally, we reported information of the gene family (Orthogroup_id) including the number of both *Bd* and total genes in each gene family (Number_Bd_genes and Total_gene_number, respectively) and molecular evolution: categorization based on origin and evolutionary history (Gene_familie_category), nucleotide substitution ratio under M0 and MA evolutionary models (M0_dN/dS and MA_2a_foreground_dN/dS, respectively).

**File S2 (separate file)**. Complete code for all analyses presented in the article.
